# CFI-1/ARID3 functions unilaterally to restrict gap junction formation in *C. elegans*

**DOI:** 10.1101/2024.04.14.589464

**Authors:** Zan Wu, Lin Pang, Mei Ding

## Abstract

Electrical coupling is vital to neural communication, facilitating synchronized activity among neurons. Despite its significance, the precise mechanisms governing the establishment of gap junction connections between specific neurons remain elusive. Here, we identified that the PVC interneuron in *Caenorhabditis elegans* forms gap junction connections with the PVR interneuron. The transcriptional regulator CFI-1/ARID3 is specifically expressed in the PVC but not PVR interneuron. Reducing *cfi-1* expression in the PVC interneuron leads to enhanced gap junction formation in the PVR neuron, while ectopic expression of *cfi-1* in the PVR neuron restores the proper level of gap junction connections in the PVC neuron, along with the normal touch response. These findings unveil the pivotal role of CFI-1/ARID3 in bidirectionally regulating the formation of gap junctions within a specific neuronal pair, shedding light on the intricate molecular mechanisms governing neuronal connectivity *in vivo*.

**SUMMARY STATEMENT:** Proper electrical coupling could be achieved by either of the partner cells; however distinctive influence on neuron circuits maybe achieved.

## Introduction

Across the nervous system, neurons mainly communicate with each other through chemical and electrical synapses (Jabeen and Thirumalai, 2018; Pereda, 2014). Chemical transmission involves the Ca^2+^-dependent release of vesicle-containing neurotransmitters from a presynaptic cell (O’Rourke et al., 2012; Südhof, 2004). In contrast, electrical synapses form intercellular channels that enable communication between the interior of two adjacent neurons (Connors and Long, 2004). Electrical coupling is prevalent during developmental stages of the nervous system (Bruzzone and Dermietzel, 2006) and plays an important role in neuronal differentiation, cell death, cell migration, synaptogenesis and neuronal circuit formation (Chalfie et al., 1985; Curtin et al., 2002; Lopresti et al., 1974; Yeh et al., 2009, Dere and Zlomuzica, 2012; Talukdar et al., 2022). In mature nervous systems, electrical synapses account for nearly 20% neuronal connections and broadly regulate neuron activity and neuronal circuits (Chalfie et al., 1985; Connors and Long, 2004; Cook et al., 2019; White et al., 1986; Zolnik and Connors, 2016). In comparison to chemical synapses, however, little is known of the mechanisms governing the formation of electrical synapses between specific pair of neurons.

Gap junctions are the biophysical substrate for the neurophysiological component of electrical synapses (Connors and Long, 2004). Connexins and innexins comprise the channel-forming proteins in vertebrate and invertebrate gap junctions, respectively (Abascal and Zardoya, 2013). Although these proteins exhibit unrelated amino acid sequences, they share remarkably similar transmembrane topologies and morphological structures. Both connexins and innexins are formed by four transmembrane-spanning domains, two extracellular and one intracellular loop, and cytoplasmic C- and N-terminal endings (Sohl et al., 2005). Usually, six or eight connexins or innexins undergo oligomerization to form a hemichannel (Stauffer et al., 1991; Oshima et al., 2016). Hemichannels in adjacent neurons appose one another to form a gap junction channel (Walker and Schafer, 2020). Ectopically expressing vertebrate connexin36 proteins could form electrical synapses and reprogram behavior in *C. elegans* (Choi et al., 2020; Rabinowitch et al., 2014). Meanwhile, innexin-containing gap junctions can function in vertebrates (Dykes et al., 2004; Phelan et al., 1998), suggesting a conserved mechanism for gap junction assembly. Recent studies further indicate that, with their powerful genetics and accessibility to live imaging, invertebrate model systems can be exploited to identify regulators of gap junction formation in mammals (Palumbos et al., 2021).

In cultured cells, overexpression of either connexins or innexins is sufficient to drive the assembly of gap junctions at the interface between random pairs of adjacent cells (Elfgang et al., 1995; Rabinowitch et al., 2014; Teubner et al., 2000). *In vivo*, however, gap junctions do not assemble between adjacent neurons that express compatible gap junction subunits (Bhattacharya et al., 2019; Fukuda, 2017; Greb et al., 2017; White et al., 1992; Yao et al., 2016). Thus, the assembly of neuron-specific electrical synapses requires additional regulatory mechanisms. Furthermore, in contrast to chemical synapses, which are primarily located at axonal boutons, electrical synapses can couple various neuronal compartments and processes (Alcami and Pereda, 2019), adding another layer of complexity to their functional impact. Gap junctions often occur as so-called gap junction plaques that contain up to thousands of gap junction channels, and presence in aggregates has been an important criterion for the identification of gap junctions (Goodenough and Paul, 2009). A functional gap junction channel is formed by two opposing hemichannels located in the plasma membrane of targeting cells. It is unclear how individual electrically coupled cells contribute to the formation of gap junctions and whether the formation of gap junctions is controlled by one side or conjointly by both sides of the partner cells.

Here, we identified that the PVC interneuron forms gap junction connections with the PVR interneuron in *C. elegans*. The transcription regulator CFI-1/ARID3 is specifically expressed in the PVC interneuron. Reducing *cfi-1* expression in PVC is sufficient to enhance gap junction formation in its partner PVR neuron. Conversely, the ectopic expression of *cfi-1* in PVR inhibits gap junction formation in PVC neuron. Beyond its rescue activity, providing *cfi-1* in PVR leads to additional behavioral consequences, indicating that the unilateral regulation of gap junction formation could exert a distinct influence on neuron circuits. This influence is likely due to the unique connectivity of each electrical coupling cell in the neuronal network.

## Results

### *cfi-1* inhibits gap junction formation in P*unc-53* expressing neurons

The gap junction protein innexin UNC-9 (Uncoordinated 9) is widely distributed in the *C. elegans* nervous system (Barnes and Hekimi, 1997). Utilizing P*unc-53* promoter-driven GFP-tagged UNC-9, we previously demonstrated its efficacy in highlighting gap junction connections established by BDU interneuron and PLM touch sensory neuron (Zhang et al., 2013). In addition to BDU, neurons expressing P*unc-53* also include PVP, PVQ, ALN, PLN, and several unidentified neurons in the tail region (Fig. 1D) (Stringham et al., 2002). Hence, gap junctions formed by those neurons could also be revealed. PVQ neurons (PVQL and PVQR), with cell bodies situated in lumbar ganglia, extend neurites anteriorly and form gap junctions to each other around the anus region. Indeed, with P*unc-53*::UNC-9::GFP marker, distinctive fluorescence puncta could be observed in anus region. Double labeling confirmed that these UNC-9::GFP puncta are indeed situated on PVQ neurons (Fig. S1A,B). In addition to the anus region, P*unc-53* driven UNC-9::GFP puncta also appear on the posterior ventral cord (Fig. 1A,B). However, prior to our study, the specific neurons responsible for forming these gap junction connections had not been unidentified.

**Fig. 1.**
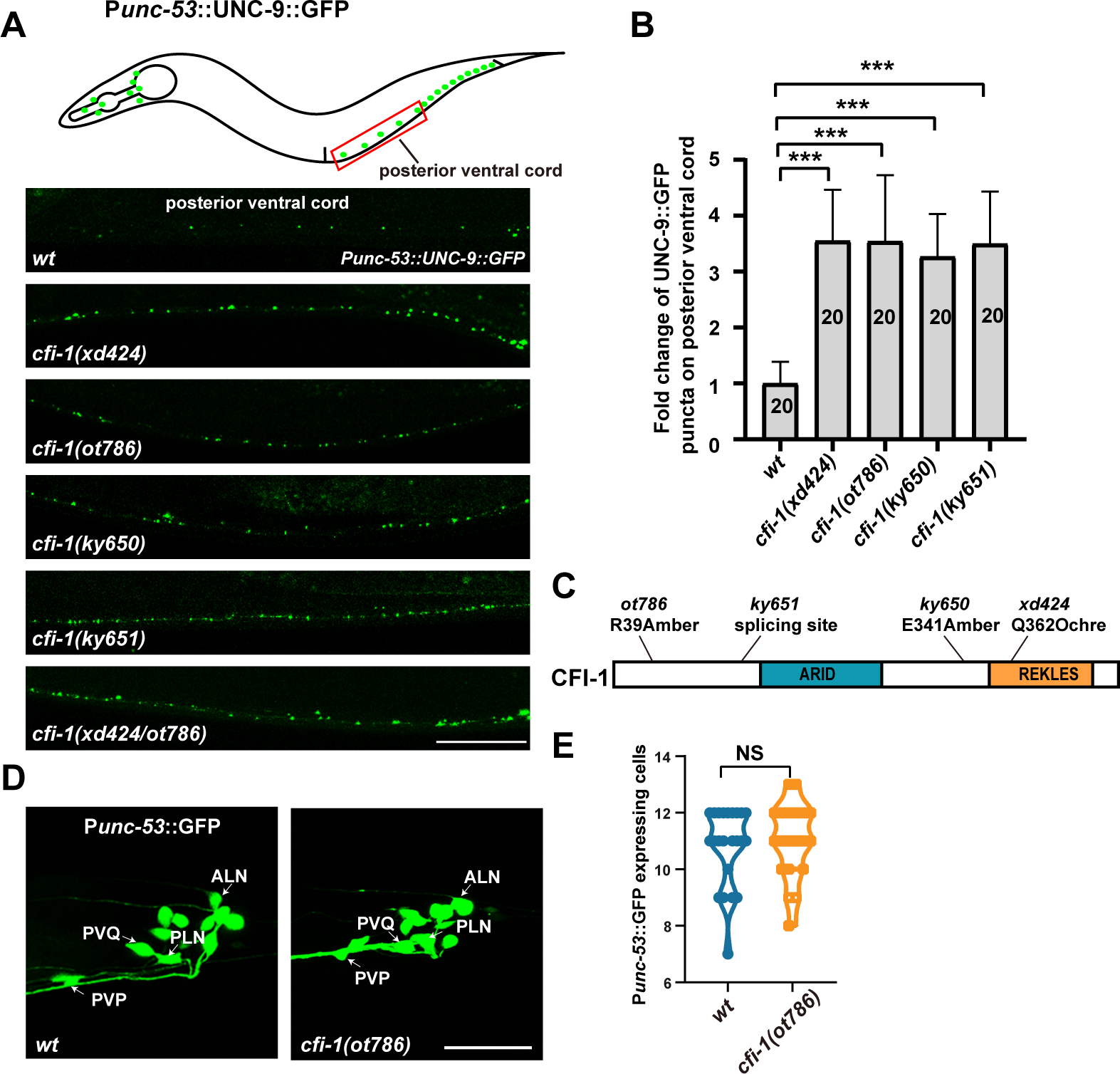
*cfi-1* inhibits gap junction formation in P*unc-53* expressing neurons. (A) Schematic drawing and the distribution P*unc-53*::UNC-9::GFP puncta (green) in wildtype (*wt*), *cfi-1 (xd424)*, *cfi-1 (ot786)*, *cfi-1 (ky651)*, *cfi-1 (ky650)*, and *cfi-1(xd424/ot786)* mutant animals. Scale bar: 25 µm. (B) The relative change of UNC-9::GFP puncta number in various genotypes. One-way ANOVA with Tukey’s multiple comparisons test was performed. Data are presented as mean ± s.d. ***, P < 0.001. N = 20 for each genotype. (C) Domain distribution of CFI-1. The mutation sites in different alleles are labeled. (D) The neurons labeled by P*unc-53* driven GFP in wildtype (*wt*) and *cfi-1 (xd424)* animals. (E) Quantification of the number of P*unc-53*::GFP expressing cells in wildtype (*wt*) and *cfi-1 (xd424)* animals. Scale bar: 25 µm. Student’s *t*-test. NS: not significant, N ≥ 20.

To explore the molecular mechanism underlying gap junction formation *in vivo*, we conducted a genetic screen to identify mutants exhibiting altered distribution of P*unc-53*::UNC-9::GFP signal. From this screen, we isolated the *xd424* mutant. In *xd424* animals, there was a noticeable increase in the number of UNC-9::GFP puncta in the posterior ventral cord region (Fig. 1A boxed region, B, and C), while the distribution of UNC-9::GFP puncta in the head and tail region remained relatively unchanged (Fig. S1A,C,D). Through genetic mapping and whole-genome sequencing, we identified a nonsense mutation (Q362Ochre) in the *cfi-1* gene in the *xd424* mutant genome (Fig. 1C). *cfi-1* encodes a worm homolog of yeast DRI and mammalian ARID3A/B proteins. All ARID family members feature a DNA-binding domain initially identified for its interaction with AT-rich DNA elements. Functioning as transcription regulators, ARID proteins have been implicated in regulating cell growth, differentiation, and development (An et al., 2010; Joseph et al., 2012; Ratliff et al., 2016). However, whether the ARID family plays a role in gap junction formation is unclear.

Other *cfi-1* alleles, including *cfi-1(ot786)*, *cfi-1(ky651)* and *cfi-1(ky650)* mutants, display increased UNC-9::GFP puncta phenotype (Fig. 1A-C) similar to *xd424* animals. Complementary test further showed that *xd424* is a *cfi-1* allele (Fig. 1A). The augmented presence of gap junction puncta in *cfi-1* mutant animals may potentially arise from an elevation in P*unc-53* expressing neurons. To test this possibility, we examined the P*unc-53*-driven GFP marker and found that the number of P*unc-53* expressing neurons is not increased by the *cfi-1(ot786)* mutation (Fig. 1D,E). This suggests that *cfi-1* regulates gap junction formation unlikely through altering the expression of P*unc-53* promoter.

### *cfi-1* could function in P*unc-53* or non-P*unc-53-*expressing neurons

Introducing a wild-type copy of *cfi-1* into P*unc-53* expressing neurons suppressed the increased gap junction puncta phenotype of *xd424* (Fig. 2A,B), indicating that loss of *cfi-1* function is responsible for the increased gap junction formation defect. Additionally, *cfi-1* could act cell-autonomously within P*unc-53* expressing neurons to regulate gap junction formation.

**Fig. 2.**
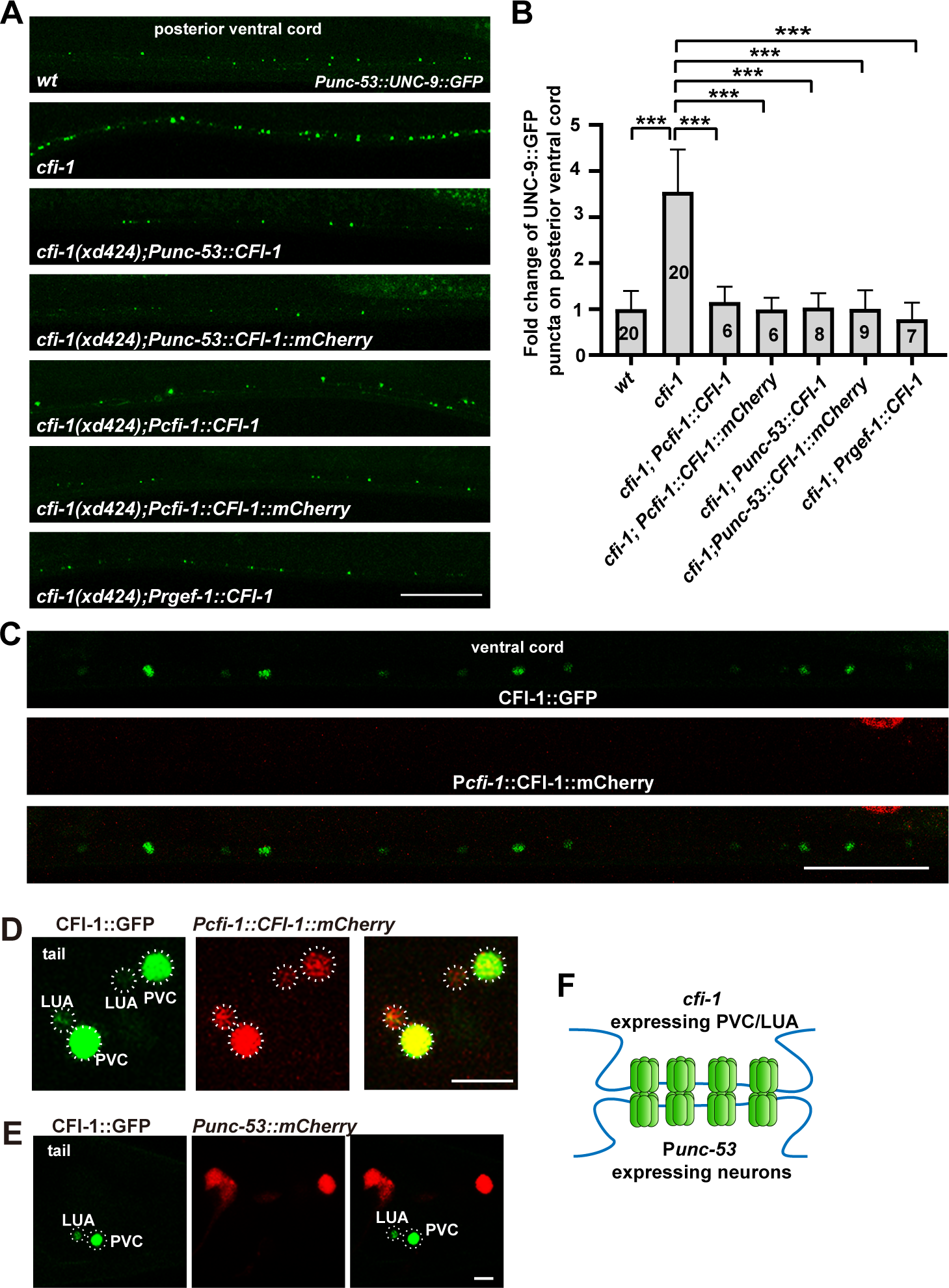
*cfi-1* could function within or outside P*unc-53* expressing neurons. (A) The ectopic UNC-9::GFP puncta (green) phenotype in *cfi-1* mutants could be rescued by P*unc-53* or P*cfi-1* promoter-driven CFI-1. Scale bar: 25 µm. (B) The relative change in UNC-9::GFP puncta number in various genotypes. One-way ANOVA with Tukey’s multiple comparisons test was performed. Data are presented as mean ± s.d. ***, P < 0.001. (C and D) Co-localization analysis of CFI-1::GFP knock-in (green) and P*cfi-1* driven CFI-1::mCherry (red) in the ventral cord (C) and tail region (D). Scale bar in (C): 25 µm. Scale bar in (D): 5 µm. (E) CFI-1::GFP knock-in (green) signal does not merge with P*unc-53* driven mCherry (red) in the tail region. Scale bar: 5 µm. (F) The *cfi-1* expressing PVC or/and LUA neurons may form gap junctions with P*unc-53* expressing neurons.

Surprisingly, however, the *cfi-1* gene appears not be expressed in P*unc-53* expressing neurons. In CRISPR/Cas9-created CFI-1::GFP knock-in (KI) line, CFI-1::GFP KI positive cells are mainly distributed in pharyngeal muscles and neurons in the head and ventral cord region (Fig. 2C,D, S2A,B) (Kerk et al., 2017). In the tail region, *cfi-1* is exclusively expressed in two PVC (PVCL and PVCR) and two LUA (LUAL and LUAR) neurons (Fig. S2A). Double labeling analysis further indicated that the nuclear localized CFI-1::GFP signal does not overlap with any P*unc-53*::mCherry expressing cells (Fig. 2E). Pan-neuronally expressing *cfi-1* could rescue the excessive gap junction phenotype of *cfi-1* mutant (Fig. S2C), while P*bnc-1* or P*unc-129* promoter-driven *cfi-1* expression in cholinergic motor neurons and/or body wall muscles in or surrounding ventral nerve cord could not suppress the increased UNC-9::GFP puncta phenotype in *cfi-1(xd424)* mutants (Fig. S2C). Intriguingly, a P*cfi-1* promoter, whose expression is limited to PVC and LUA neurons in the tail region (Fig. S3A), driven wild-type *cfi-1*, could significantly suppress the increased UNC-9::GFP puncta phenotype of *cfi-1* mutants (Fig. 2A). Hence, endogenous *cfi-1* may function in LUA or/and PVC to regulate gap junction formation in P*unc-53* expressing neurons.

### PVC neurons are gap junction partners for P*unc-53* expressing neurons

Providing *cfi-1* in non-P*unc-53* expressing LUA and PVC neurons successfully rescued the *cfi-1* mutant phenotype, indicating a potential cell-non-autonomous function of *cfi-1*. How does this cell-non-autonomous rescue occur? We hypothesized that PVC or/and LUA might form gap junction connections with certain P*unc-53* expressing neurons (Fig. 2F) and the loss of *cfi-1* function in PVCs or/and LUAs resulted in defects in gap junction formation in the partner cells of PVCs or/and LUAs, as revealed by the P*unc-53*::UNC-9::GFP marker. If this hypothesis holds, with which P*unc-53* expressing neurons the PVC or/and LUA form gap junctions? To address this question, we firstly examined whether LUA or PVC neurons form gap junctions on the posterior ventral cord. LUA neurons extend relatively short neurites anteriorly, terminating at the anus region (lumbar commissures), and do not reach the posterior ventral cord region where UNC-9::GFP puncta are observed. In contrast, PVC neurites extend to the nerve ring region in the head (White et al., 1986). Based upon, we suspected that the UNC-9::GFP puncta on the posterior ventral cord are more likely to form on partner cells of PVC neurons rather than LUA neurons. Indeed, when we co-expressed P*unc-53*::UNC-9::GFP with the PVC-specific marker (P*nmr-1* promoter driven mCherry) (Shaham and Bargmann, 2003), we found that the UNC-9::GFP puncta on the posterior ventral cord are align with PVC neuronal processes (Fig. 3A,4G).

**Fig. 3.**
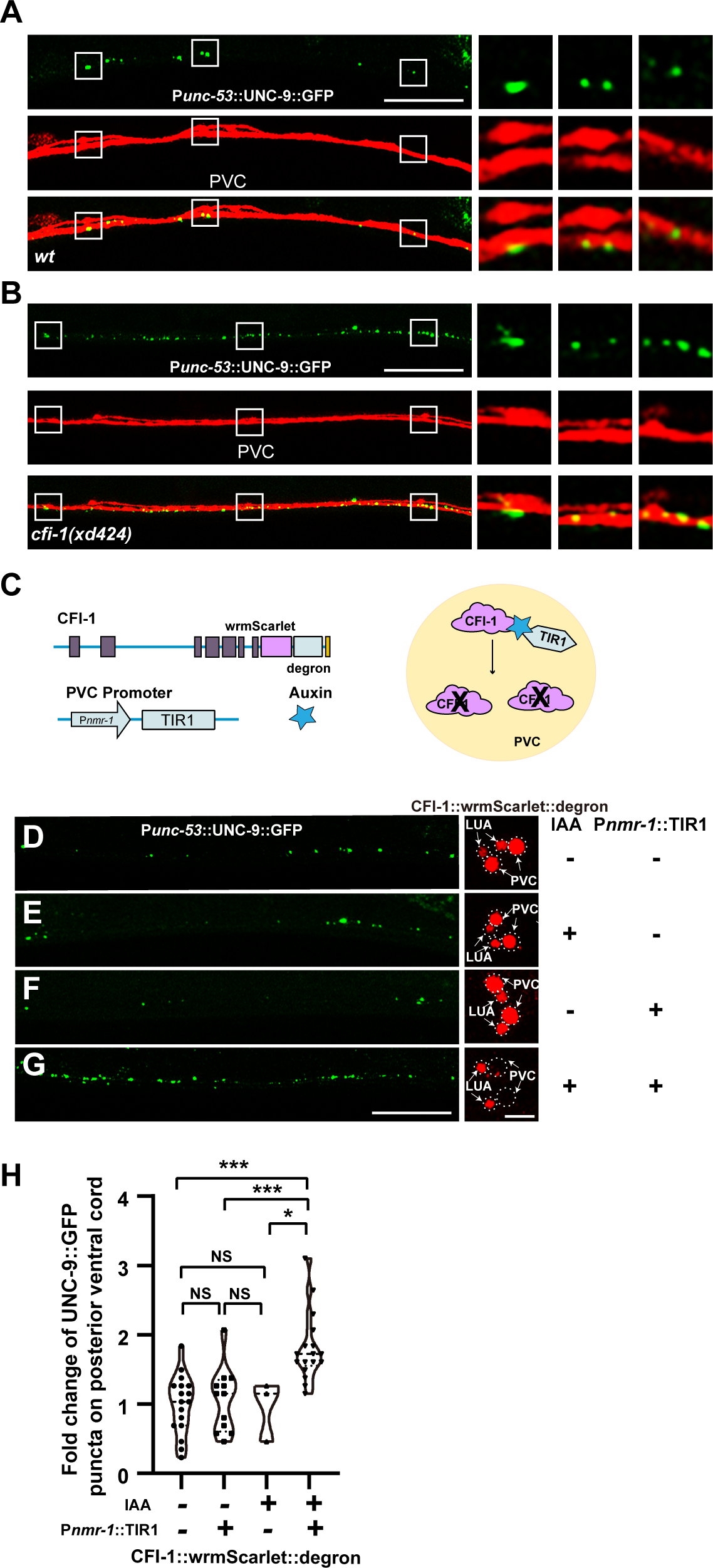
*cfi-1* functions in PVC neurons. (A and B) Gap junctions formed by P*unc-53* expressing neurons (green, labeled by P*unc-53*::UNC-9::GFP) are situated on PVC neurons (red) in wild type (A) and *cfi-1(xd424)* mutant (B) animals. The single layers of the Z-stacks of boxed regions are enlarged on the right. (C) Schematic drawing of CFI-1 AID in PVC neurons. (D-H) Expression of CFI-1::wrmScarlet::degron (red) and UNC-9::GFP puncta (green) driven by P*unc-53* with or without P*nmr-1*::TIR1 or IAA. Scale bar: 25 µm and 5 µm. (H) The relative change in UNC-9::GFP puncta number in various treatments. One-way ANOVA with Tukey’s multiple comparisons test was performed. Data are presented as mean ± s.d. ***, P < 0.001. *, P < 0.05. NS: not significant.

**Fig. 4.**
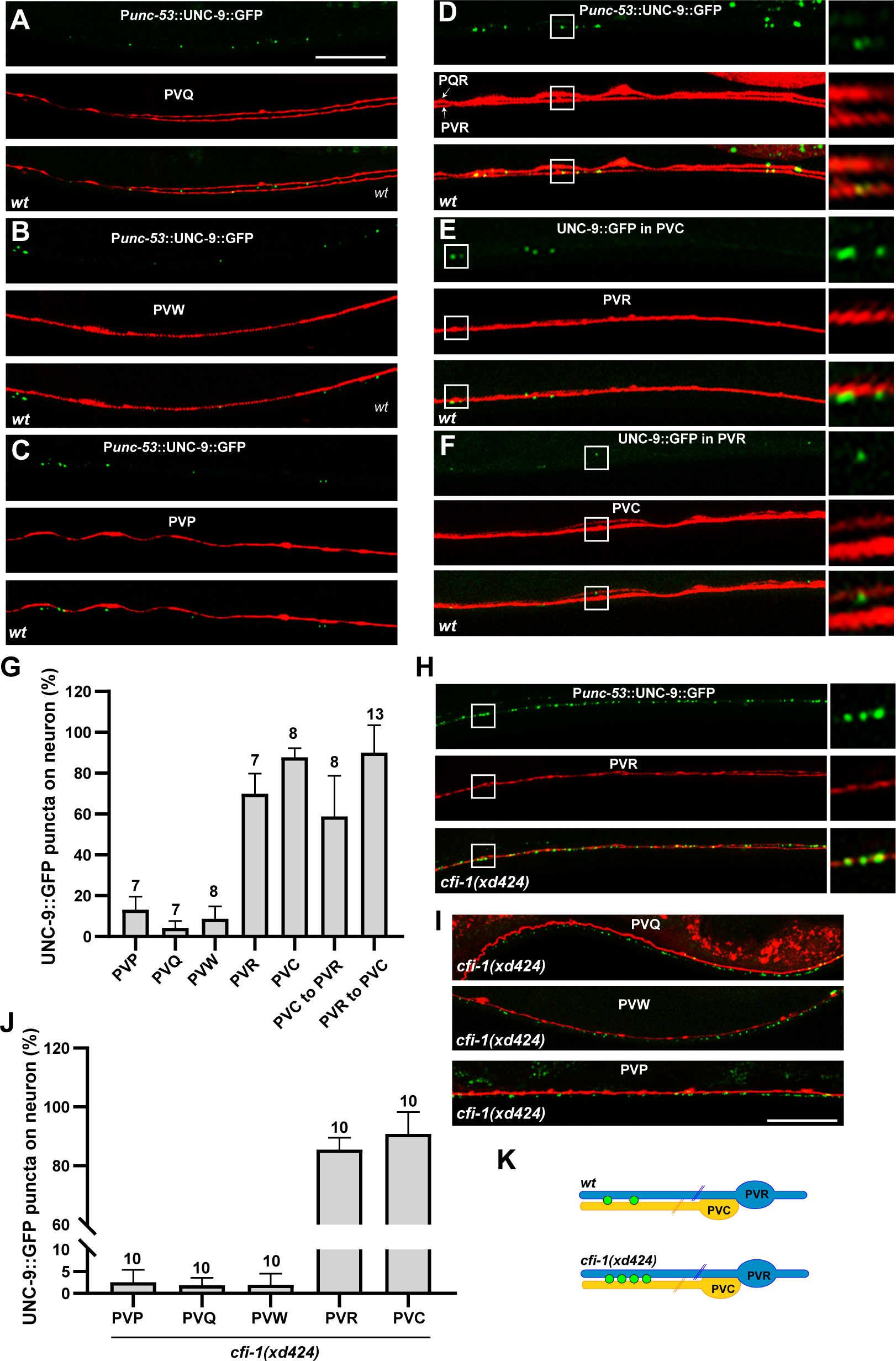
PVC and PVR are gap junction partner cells. (A-C) The P*unc-53*::UNC-9::GFP puncta (green) are not present on PVQ (red) (A), PVW (red) (B) or PVP (red) (C). (D) The P*unc-53*::UNC-9::GFP puncta (green) are present on PVR (red). The single layers of the Z-stacks of boxed regions are enlarged on the right. (E) The P*nmr-1*::UNC-9::GFP puncta (green) (from PVC) are situated on PVR (red) neurons. The single layers of the Z-stacks of boxed regions are enlarged on the right. (F) The P*flp-10*::UNC-9::GFP puncta (green) (from PVR) are situated on PVC (red) neurons. The single layers of the Z-stacks of boxed regions are enlarged on the right. Scale bar: 25 µm. (G) Quantification of the percentage of UNC-9::GFP puncta on various types of neurons. Data are presented as mean ± s.d. (H and I) The ectopic P*unc-53*::UNC-9::GFP puncta (green) observed in *cfi-1(xd424)* mutant animas are formed by PVR (red) (H), but not by PVQ (red), PVW (red) or PVP (red) (I). The single layers of the Z-stacks of boxed regions are enlarged on the right. Scale bar: 25 µm. (J) Quantification of the percentage of UNC-9::GFP puncta on various types of neurons in *cfi-1(xd424)* mutant animals. Data are presented as mean ± s.d. (K) Schematic drawing of gap junction connections between PVC and PVR neurons in wild type (*wt*) and *cfi-1(xd424)* mutants.

We further tested whether *cfi-1* function in PVC neurons is required for the formation of gap junctions in P*unc-53* expressing neurons. To address this, we utilized the auxin-inducible degradation (AID) system (Zhang et al., 2015). In the *C. elegans* genome, we integrated the wrmScarlet::degron DNA sequence into the 3’-terminal end of the *cfi-1* coding region. Subsequently, we expressed the TIR1 E3 ligase in PVC neurons using the P*nmr-1* promoter. Upon exposure to auxin, the wrmScarlet signal was selectively reduced in PVC neurons but remained unchanged in LUA neurons, indicating the specific depletion of CFI-1 protein in PVC neurons. When *cfi-1* function was specifically reduced in PVCs, we observed a substantial increase in the number of P*unc-53*::UNC-9::GFP puncta (Fig. 3C-H). Together, above data collectively suggest that *cfi-1*-expressing PVC neurons form gap junctions with P*unc-53* expressing neurons, and *cfi-1* function in PVCs is crucial for suppressing gap junction formation in P*unc-53* expressing neurons.

### PVR is the gap junction partner cell for PVC neurons

Next, we went to identify with which P*unc-53* expressing neurons the PVCs form gap junction. The known P*unc-53* expressing neurons include ALNs, PLNs, PVPs, and PVQs. Co-labeling experiments with markers, including P*lad-2*::mCherry (ALN and PLN), P*odr-2b*::mCherry (PVP), and P*sra-6*::mCherry (PVQ), confirmed the expression of P*unc-53* in these neurons (Chou et al., 2000; Troemel et al., 1995; Wang et al., 2008;) (Fig. S3B-D). Additionally, the P*unc-53*::GFP signal was observed in unidentified neurons in the tail region. Through further co-labeling experiments, we identified these unknown P*unc-53* expressing neurons as PVWs (labeled by P*flp-7*) (Kim and Li, 2004), PVR (labeled by P*flp-10*), and PQR (also labeled by P*flp-10*) (Kim and Li, 2004) (Fig. S3E,F), but not PLM (labeled by P*mec-7*::mCherry) (Mitani et al., 1993) or PVN (labeled by P*pdf-1*::mCherry) (Barrios et al., 2012) neurons (Fig. S3G,H).

Among the P*unc-53* expressing neurons, ALN and PLN neuronal processes are positioned laterally, away from the ventral cord (Fig. S3B), making it unlikely for them to form gap junctions with PVCs. PVW neurons form gap junctions with PVC neurons but in the distal anterior region of the ventral cord (White et al., 1986). Apart from that, there is limited information about which P*unc-53* expressing neurons may form gap junctions on PVCs. Therefore, to identify the partner cells of PVCs in gap junction formation, we further employed the P*unc-53*::UNC-9::GFP marker and examined the distribution of GFP puncta on PVP, PVQ, PVW, PVR, or PQR neurons. Our survey revealed that P*unc-53*-driven UNC-9::GFP puncta are absent on PVQs (Fig. 4A,G), PVWs (Fig. 4B,G), or PVPs (Fig. 4C,G). In contrast, on the posterior ventral cord, the UNC-9::GFP puncta are exclusively localized on the PVR neuron (Fig. 4D,G). We further co-labeled the PVR neuron and PVC gap junctions and found that PVCs indeed form gap junctions with the PVR neuronal process (Fig. 4E, G). Conversely, PVR gap junctions are specifically situated on PVC neurons (Fig. 4F, G). Taken together, PVCs and PVRs form gap junctions to each other (Fig. 4K).

Does *cfi-1* influence the gap junction formation between PVC and PVR neurons? Co-labelling experiments with the PVR neuron revealed that the excessive P*unc-53*::UNC-9::GFP puncta in *cfi-1* mutants were localized on PVR (Fig. 4H), rather than on PVQ, PVP, or PVW neurons (Fig. 4I, J, and K). Conversely, when examining PVC neurons, we found that the surplus UNC-9::GFP puncta were positioned on PVC neurons (Fig. 3B). Therefore, *cfi-1* specifically regulates the formation of gap junctions between PVC and PVR neuronal pairs.

### *cfi-1* specifically regulates the gap junction formation of PVC-PVR pair

So far, we had been utilizing promoter-driven UNC-9::GFP to examine the spatial arrangement of gap junction connections in *C. elegans* neurons. To address potential concerns regarding overexpression, we further employed the Native and Tissue-specific Fluorescence (NATF) technique. NATF combines CRISPR/Cas9-mediated genome editing with the split-GFP method, facilitating the specific labeling of proteins within tissues at their endogenous levels (He et al., 2019). Basically, we divided the GFP molecule into two fragments, GFP1-10 and GFP11. The DNA sequence encoding the GFP11 fragment was inserted into the 3’-end of the *unc-9* gene via the CRISPR/Cas9 method. In UNC-9::GFP11 knock-in worms, no green fluorescence signal was detectable. We then introduced the tissue-specific expressing promoter-driven GFP1-10 fragment into UNC-9::GFP11 knock-in worms. This allowed for the characterization of endogenous gap junctions within specific tissues or cells (Fig. 5A,B). In comparison to P*unc-53*-driven UNC-9::GFP, the signal from NATF-mediated P*unc-53*-driven UNC-9::split-GFP was dimmer, with fewer visible UNC-9::split-GFP puncta (Fig. S4A,B). Nevertheless, consistent with P*unc-53*-driven UNC-9::GFP, UNC-9::split-GFP revealed the presence of PVQ gap junctions in the anus region (Fig. S1B, S4A,B boxed region). Similarly, puncta of UNC-9::split-GFP were observable in the posterior ventral cord region, indicating a similar distribution pattern of gap junctions *in vivo* between P*unc-53*::UNC-9::GFP and P*unc-53*::UNC-9::split-GFP.

**Fig. 5.**
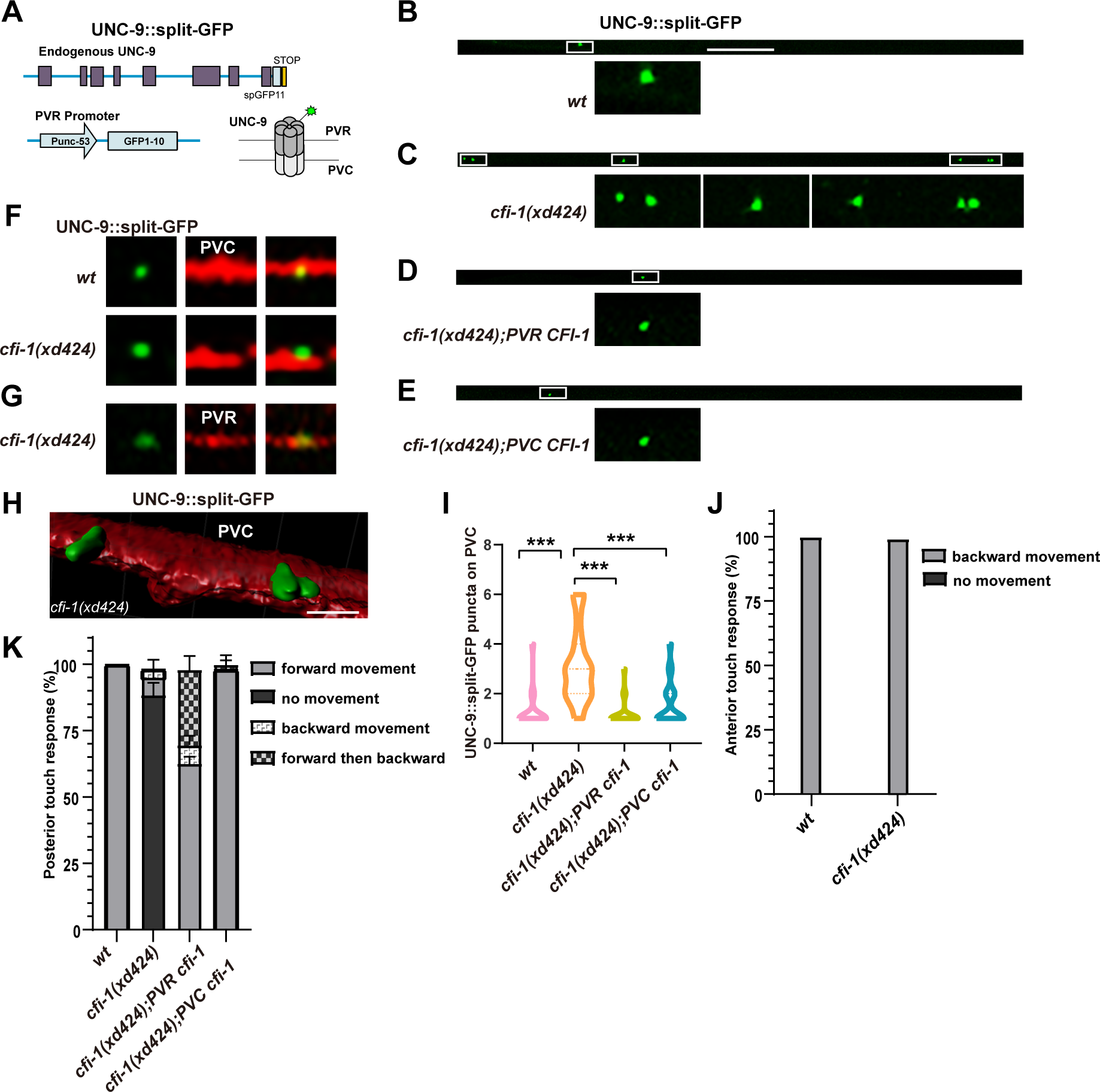
*cfi-1* affects gap junction formation between PVC and PVR neurons. (A) Schematic drawing of endogenous gap junctions labeling by NATF approach. (B-E) The UNC-9::split-GFP (green) puncta formed by PVR neurons in wild type (*wt*) (B), *cfi-1(xd424)* (C), *cfi-1(xd424);PVR CFI-1* (D), or *cfi-1(xd424);PVC CFI-1* (E) animals. Scale bar: 25 µm. (F) The endogenous gap junctions (labeled by UNC-9::split-GFP) formed by PVR neuron are situated on PVC (red) neurons in both wild type and *cfi-1(xd424)* mutant animals. (G) The ectopic gap junctions (labeled by UNC-9::split-GFP) formed by PVR neuron are indeed found on PVR (red) neurons in *cfi-1(xd424)* mutant animals. (H) The 3-D reconstruction of endogenous gap junctions (labeled by UNC-9::split-GFP) formed by PVR neurons on PVC (red) neurons in *cfi-1(xd424)* mutant animals. Scar bar: 2 µm. (I) Quantification of the number of endogenous gap junctions formed by PVR neuron in various genotypes. One-way ANOVA with Tukey’s multiple comparisons test was performed. N ≥ 21 for each genotype. ***, P < 0.001. (J) The anterior touch response in wild type and *cfi-1(xd424)* mutant animals. (K) The posterior touch response in various genotypes. Data are presented as mean ± s.d. N ≥ 18.

Upon introducing the PVC marker into the P*unc-53*::UNC-9::split-GFP expression line, we found that the UNC-9::split-GFP puncta on the posterior ventral cord are specifically located on PVC neurons (Fig. 5F). This observation supports the idea that PVCs form gap junctions onto P*unc-53* expressing PVR neuron. Notably, aside from PVC-PVR gap junctions, no other discernible UNC-9::split-GFP puncta were evident in the posterior ventral cord region. In *cfi-1* mutant animals, the number of UNC-9::split-GFP puncta along the posterior ventral cord is increased (Fig. 5C,I). Through double staining, we confirmed that the surplus UNC-9::split-GFP puncta in *cfi-1* mutants were formed between PVC (Fig. 5F) and PVR neurons (Fig. 5G). Additionally, we generated a 3-D reconstruction image, illustrating that the green P*unc-5*3::UNC-9::split-GFP puncta are embedded within PVC neurons (Fig. 5H). Could *cfi-1* function in either PVC or PVR to regulate the formation of endogenous PVC-PVR gap junctions? Utilizing the UNC-9::split-GFP marker, we found that introducing wild type *cfi-1* gene into PVR or PVC could lead to significant rescue effect in *cfi-1* mutants (Fig. 5D,E,I). Together, these findings further support the notion that *cfi-1* operates bidirectionally in either PVC or PVR to regulate gap junction formation between PVC and PVR neurons (Fig. 4K).

Given the broad expression of *cfi-1* along the ventral cord, we further investigated its impact on endogenous gap junction formation in motor neurons through the NATF approach. We found that *cfi-1* mutation does not impede the formation of gap junction connections in DD/VD neurons (Fig. S4C,D). Thus, *cfi-1* may selectively regulate gap junction formation in the PVC-PVR neuronal pair.

### *cfi-1* could regulate touch response from either PVR or PVC neuron

Electrical coupling through gap junctions dynamically influences neuronal circuitry and behavioral outcomes. In *C. elegans*, PVC serves as a major command interneuron for mediating forward locomotion triggered by posterior touch (Fig. S5A) (Chalfie et al., 1985). PVR forms gap junctions with touch sensing neurons ALM and PLM (White et al., 1986), which are responsible for receiving anterior and posterior touch stimuli, respectively (Chalfie et al., 1985). Here, we discovered that PVR also forms gap junctions with PVC interneurons (Fig. S5A), and mutations in *cfi-1* lead to excessive gap junction formation rather specifically between PVC and PVR neurons (Fig. S5B). Notably, *cfi-1* mutant animals have been shown to exhibit defects in touch response (Shaham and Bargmann, 2003). Specifically, when gentle touch stimuli were applied around the posterior region, the anterior movement of *cfi-1* mutant animals was inhibited (Fig. 5K), while touch stimuli were applied around the anterior region resulted in posterior locomotion similar to wild type (Fig. 5J). This observation is consistent with the role of *cfi-1* in restricting electrical coupling between PVC and PVR neurons. Introduction of wild type *cfi-1* into PVC neurons successfully rescued the posterior touch response defect in *cfi-1* mutant animals, highlighting the critical role of PVCs in posterior touch response (Fig. 5K and Fig. S5A). Excessive gap junction formation between PVC and PVR neurons could be inhibited by expressing *cfi-1* in PVR neuron. Remarkably, introducing wild type *cfi-1* into PVR neurons also suppressed the posterior touch response defect of *cfi-1* mutants (Fig. 5K). We noticed that when *cfi-1* was expressed only in PVR but not in PVC, some animals exhibited abnormal backward movement upon posterior touch (Fig. 5K). Given the distinct connectivity patterns of PVC and PVR, it is possible that individual partner cells of the electrically coupled neuron pair may have distinct outputs to the touch circuit (Fig. S5C and Discussion). Nevertheless, the rescuing activity of *cfi-1* in either PVC or PVR supports the notion that unilateral bidirectional regulation of gap junction formation could profoundly influence neuron function and subsequently, the behavioral outcomes controlled by relevant neural circuits.

## Discussion

The nearly completed connectome of the simple worm nervous system offers a distinctive and pivotal model for elucidating the principles governing gap junction formation *in vivo*. Here, we revealed that the gap junction formation can be orchestrated by either side of the partner cells. Given that a functional gap junction is constituted by hemichannels originating from opposing cells, this unilateral bidirectional regulatory mechanism could exert intricate and profound effects on the neuronal network.

The *C. elegans* genome contains a single ARID3 ortholog CFI-1, whereas mammals contain three ARID3 subfamily members (ARID3A, ARID3B and ARID3C) (Kortschak RD, 2000; Shaham and Bargmann, 2002; Wilsker et al., 2005). Previous studies indicated that ARID3 members participate in embryonic patterning, cell lineage, cell cycle, apoptosis, stem cell differentiation, tumorigenesis, and so on (An et al., 2010; Herrscher et al., 1995; Joseph et al., 2012; Numata et al., 1999; Ratliff et al., 2014; Ratliff et al., 2016; Roy et al., 2014; Shandala T et al., 1999; Valentine SA et al., 1998; Webb et al., 2011;). In *C. elegans*, *cfi-1* was originally cloned based upon its role in the development of neuronal diversity (Shaham and Bargmann, 2002). Through the generation of an *in vivo* binding map on the *C. elegans* genome, Li et al. recently identified that the majority of CFI-1 targeting genes encode markers associated with neuronal terminal differentiation (Li et al., 2023). Therefore, CFI-1 may act a terminal selector, orchestrating the terminal differentiation of distinct neuron types. In line with this study, we found a tight correlation between *cfi-1* expression in PVCs and gap junction formation in the PVC-PVR neuronal pair. Notably, electrical coupling emerges as a crucial feature in neuronal terminal differentiation. Here, *cfi-1* does not appear to regulate the specificity of gap junction formation; instead, it modulates the quantity or abundance of gap junctions formed between appropriate partner cells. Thus, terminal selectors may also exert control over the electrical coupling strength of specific neuronal subtypes

As a transcription regulator, how could CFI-1/ARID3 achieve inhibition of gap junction formation from one side of the partner cells? In PVC-PVR pair, endogenous *cfi-1* exhibits expression solely in PVC neurons, not in the PVR neuron. Within the PVCs, CFI-1 may directly modulate the gene expression of gap junction proteins, thereby exerting control over the quantity or abundance of gap junction connections. The whole genome binding map did not identify innexin *unc-9* as a direct target gene of CFI-1 (Li et al., 2023). However, given the limited number of neurons (confined to PVC and PVR) in which gap junction formation is influenced by *cfi-1*, it is difficult to rule out the possibility that *cfi-1* could directly regulate the gene expression of UNC-9 or other gap junction proteins. Alternatively, CFI-1 is involved in the transcription regulation of genes associated with the stability of gap junctions. While numerous proteins in cultured cells have been identified as regulators of gap junction homeostasis, few studies have investigated how the abundance of gap junction connections is determined in an intact nervous system. Recent research has demonstrated that the conserved CASPR family nematode protein NLR-1 anchors the F-actin at the gap junction plaque, and plays a critical role in gap junction assembling (Meng and Yan, 2020). The cAMP-dependent signaling pathway regulates the trafficking of gap junction proteins, thereby controlling the subcellular localization of *C. elegans* gap junction connections (Palumbos et al., 2021). Neurobeachin, a conserved protein with multiple protein binding domains, localizes to electrical synapses and regulates the robust localization of neuronal connexins post-synaptically (Martin et al., 2023; Miller et al., 2015). However, the specific mechanistic involvement of CFI-1 in modulating the trafficking or stability of gap junction proteins within established cellular pathways, leading to the inhibition of gap junction formation, remains elusive. How does the formation of gap junctions in PVC neurons influence the corresponding process in PVR neurons? We suspect that CFI-1 may regulate gap junction formation in opposing cells indirectly by modulating the transcription of unidentified extracellular signaling molecules. Alternatively, the formation or connexin/innexin hemichannel clusters in one cell may directly impact the clustering of connexin/innexin hemichannels in another cell. This direct effect likely arises from the stability of gap junctions facilitated by interaction between juxtaposed hemichannels (Sosinsky and Nicholson, 2005). It is noteworthy, however, that the expression of compatible gap junction subunits does not guarantee the assembly of gap junction connections between adjacent neurons *in vivo* (Bhattacharya et al., 2019; Fukuda, 2017; Greb et al., 2017; White et al., 1992; Yao et al., 2016). Thus, while the formation or connexin/innexin hemichannel clusters in one cell may influence the clustering of connexin/innexin hemichannels in another cell, the assembly of neuron-specific electrical synapses likely involves additional regulatory mechanisms.

*cfi-1* is expressed in diverse motor neurons within the ventral cord. However, the overall distribution of gap junctions on ventral cord motor neurons does not appear to be affected by *cfi-1* mutations. This highlights the cell-context dependent nature of *cfi-1* in regulating gap junctions. Consistent with this observation, CFI-1 engages in collaborative interactions with multiple transcription regulators to influence the differentiation of distinct neuronal cell types. In partnership with the POU homeodomain transcription factor UNC-86, CFI-1 directly activates terminal differentiation genes specific to IL2 neurons (Shaham and Bargmann, 2002). Concurrently, CFI-1 collaborates with two distinct homeodomain proteins, UNC-42/Prop-1 like and CEH-14/LIM, to govern the terminal differentiation processes of AVD and PVC interneurons, respectively (Glenwinkel L et al, 2021; Berghoff EG et al, 2021). The transcription factor UNC-3, an ortholog of human EBF1, EBF2, and EBF4, regulates the gene expression of *cfi-1* in motor neurons (Kerk et al., 2017). Within ventral nerve cord motor neurons, CFI-1 functions as a repressor of the glutamate receptor gene *glr-4*/GRIK4. The *glr-4* gene undergoes positive regulatory control from three conserved transcription factors, namely UNC-3/Ebf, LIN-39/Hox4-5, and MAB-5/Hox6-8, while experiencing negative regulation from CFI-1 (Kerk et al., 2017). Therefore, CFI-1 likely collaborates with distinct sets of transcription regulators in various neuronal subsets, and its functional requirement in gap junction formation may only be evident in selective neuronal populations.

The touch response could be restored when placing *cfi-1* in its non-native expressing cell, PVR, indicating that the PVC functional deficiency in *cfi-1* mutants is likely due to over-coupling between PVC and PVR neurons. Intriguingly, expressing *cfi-1* in PVR but not PVC neurons results in abnormal backward movement upon posterior touch, suggesting that the electrically coupled neuron pair is not an inseparable unit. Rather, the entire and unique connectivity of a given neuron determines its distinctive function. So, how can we explain the abnormal backward movement? In a simple model (Fig. S5C), we might suggest that when a wild-type copy of *cfi-1* is introduced into PVR, the electrical coupling between PVR and PVC is selectively reduced. Consequently, the electrical coupling between PVC and AVA may be strengthened, allowing more mechanical signal to flow into DA/AS motor neurons, thus causing the backward movement. Of course, one could explore different neuronal connections to achieve a similar behavioral outcome by altering the activity of other neurons. Our current understanding of gap junction formation is far from complete. The complex interactions among diverse terminal selectors suggest that there will likely be significant divergence in the regulatory mechanisms governing gap junction formation in individual neurons. To add further complexity, most neurons usually express more than one gap junction gene (Evans and Martin, 2002). Additionally, one side of an electrical synapse is not necessarily considered the mirror of the other (Rash et al., 2013). Therefore, determining whether *cfi-1*-mediated PVR-PVC gap junction formation would influence gap junctions between other neurons and PVC or PVR is rather challenging.

Electrical coupling could occur dendro-dendritically, somato-somatically, or between axons (Alcami and Pereda, 2019). In PVC and PVR neurons, gap junctions form in the middle of their anterior neuronal processes. The excess gap junction connections resulting from *cfi-1* loss-of-function also occur in this region. What signifies this particular region? In the ventral cord, PVC neurons receive synaptic input from various neurons including AVA, PHB, PHC, VA12, LUA, PVM, PVN, DVA and PVD, while primarily sending synaptic output to motor neurons such as VBs and DBs. Notably, PVD and DVA neurons are presynaptic to both forward and backward interneurons, contributing to both the anterior and posterior touch circuits. Conversely, PVR in the ventral cord receives synaptic input from PVM, DVA and PVM, and through gap junctions is coupled with DVA and PLM. Along with ALM and PLM, AVM neurons sense gentle mechanical stimuli to the body and provide input to command interneurons. This positioning of PVC-PVR gap junctions likely facilitates sensory-motor integration. However, how this specific subcellular localization of gap junctions is determined during development is completely unknown. Similarly, the unique roles of each neuron in specifying the location and abundance of gap junctions within the PVC-PVR pair are not yet understood. Our studies have only scratched the surface of the intricate regulatory mechanisms governing electrical synapse formation *in vivo*. Further investigation is needed to fully elucidate how characteristic electrical coupling is achieved within specific neuronal pairs in the nervous system.

## MATERIALS AND METHODS

### Worm strains and genetics

*C. elegans* strain maintenance and genetic manipulation were performed under standard conditions as described (Brenner, 1974). Mutants and transgenic fluorescence reporters used in this study are listed here: LGI, *cfi-1(xd424)*, c*fi-1(ot786)*, *cfi-1(ky650)*, *cfi-1(ky651)*, *xdKi79* (CFI-1::GFP KI), *xdKi83* (CFI-1::wrmScarlet::degron KI). LGII, xdIs174 (P*unc-53*::UNC-9::GFP, P*myo-2*::RFP), LGX, *xdKi13* (UNC-9::GFP11 KI). Additional transgenic lines are: *xdEx276* (P*unc-53*::GFP, P*odr-1*::RFP), *xdEx2366* (P*cfi-1*::GFP, P*odr-1::RFP*), *xdEx2373* (P*unc-53*::CFI-1, P*odr-1*::GFP), *xdEx2383* (P*unc-53*::CFI-1::mCherry, P*odr-1*::GFP), *xdEx2386* (P*unc-53*::GFP, P*nmr-1*::mCherry, P*odr-1*::GFP), *xdEx2442* (P*unc-53*::GFP, Pf*lp-7*::mCherry, P*odr-1*::GFP), *xdEx2445* (P*unc-53*::GFP, P*flp-10*::mCherry, P*odr-1*::GFP), *xdEx2448* (P*unc-53*::GFP, P*lad-2*::mCherry, P*odr-1*::GFP), *xdEx2462* (P*unc-53*::GFP, P*sra-6*::mCherry, P*odr-1*::GFP), *xdEx2465* (P*unc-53*::GFP, P*odr-2 2b*::mCherry, P*odr-1*::GFP), *xdEx2489* (P*unc-53*::GFP, P*cfi-1*::MYR::mCherry, P*odr-1*::GFP), *xdEx2495* (P*unc-53*::CFI-1, P*odr-1*::GFP), *xdEx2509* (P*unc-53*::GFP1-10, P*odr-1*::RFP), *xdEx2837* (P*cfi-1*::CFI-1::mCherry, P*odr-1*::GFP), *xdEx2915* (P*unc-53*::mCherry, P*odr-1*::RFP), *xdEx2918* (P*cfi-1::CFI-1*, P*odr-1*::GFP), *xdEx3051* (P*nmr-1*::mCherry; P*myo-2*::RFP), *xdEx3071* (P*flp-7*::mCherry; P*odr-1*::GFP), *xdEx3072* (P*odr-2 2b*::mCherry; P*odr-1*::GFP), *xdEx3073* (P*flp-10*::mCherry; P*nmr-1*::UNC-9::GFP; P*odr-1*::RFP), *xdEx3076* (P*nmr-1*::mCherry; P*flp-10*::UNC-9::GFP; P*odr-1*::RFP), *xdEx3079* (P*flp-10*::mCherry; P*odr*-1::GFP), *xdEx3080* (P*nmr-1*::mCherry; P*odr-1*::GFP), *xdEx3081* (P*sra-6*::mCherry; P*odr-1*::GFP), and *xdEx3117* (P*nmr-1*::TIR1; P*odr-1*::GFP).

The *cfi-1(xd424)* mutant was isolated from *xdIs174* animals treated with ethylmethane sulfonate (EMS). 10,000 mutagenized haploid genomes were screened and total sixteen mutations were isolated from this screen.

### DNA constructs and transgenes

DNA fragments were inserted into the pSM, ΔpSM, pPD95.77 or pPD95.75 vector using standard procedures. For *cfi-1* tissue specific rescue experiments, DNA constructs were injected into young adult animals at a concentration of 5ng/µL concentration. For split GFP labeling, DNA constructs containing the GFP1-10 fragment were injected into young adults animals at a concentration of 1 ng/µL. To label various neurons, the corresponding DNA constructs were injected into worms at a concentration of 50ng/µL. The co-injection marker was P*myo-2*::RFP, P*odr-1*::GFP, or P*ord-1*::RFP injected at a concentration of 50ng/µL. Integrated strains were obtained using the trimethylpsoralen/ultraviolet (TMP/UV) method (Gengyo-Ando et al., 2000).

### CRISPR/Cas9-mediated gene editing

The knock-in strains are generated by CRISPR/Cas9-mediated genome editing, and they were constructed by SunyBiotech and verified with PCR and sequencing. The sequences of sgRNA and corresponding primers are listed here. To generate the *xdKi13 (UNC-9::GFP11 KI)*, the sgRNAs (CGTATGGTTGCAACTCACGCCGG and CCGGAGAACTACCCTGTTACGAG) and primers (forward primer GGTGTTTTCCTACTTCGTATGGTTG and reverse primer GACGACTACACCCATTGACGAC) were used. To generate *xdKi79 (CFI-1::GFP KI)* and *xdKi83 (CFI-1::wrmScarlet::degron KI)*, the sgRNAs (TCAGTATCAATGGAAATCAACGG and ATCAACGGAATCACCTATCAAGG) and primers(F/s1: ATGGTGCATCGAGTATGAGGA, R/s2: CAAATTGCGATCACCGAGA, xdKi79-mid-F/s3: ATGCCCGAAGGTTATGTACAGG, xdKi83-mid-F/s3: CAGCCGACATCCCAGACTACTA, and xdKi83-mid-R/s4: TTGAAGTCGGCGAGGTAA) were used.

### Microscopy and image acquisition

Animals were placed on 2.5% agar pads in M9 buffer with 1.4% 1-phenoxy-2-propanol. Fluorescence images of nematodes were captured using the Leica SP8 confocal microscope. The photographs were taken at the young adult stage unless specifically indicated. Confocal stacks were projected into a single image. The number of UNC-9::GFP puncta with the gray value of ≥ 40 was counted. For relative UNC-9::GFP distribution on each neurons is defined as the GFP puncta number at the mCherry region on single layer of image divided by total GFP puncta number along the ventral nerve cord region. The posterior ventral cord region is defined as extending from 50 µm posterior to the vulval region to 190 µm posterior to it. The tail region is characterized as ranging from 20 µm anterior to the anus region to 120 µm anterior to it. Straightened ventral cord images were obtained using ImageJ. The 3-D reconstruction of the co-localization analysis between P*unc-53*::GFP1-10; UNC-9::GFP11 and PVC neuron marker was performed using Imaris software.

### Statistical analysis

To compare multiple groups, one-way ANOVA was used with an appropriate multiple comparisons post hoc test (the test used is stated in each figure legend). *P < 0.05; **P < 0.01; ***P < 0.001; NS, not significant.

### AID assay and auxin treatment

Temporally controlled protein degradation depletion using the AID (auxin-inducible degradation) system was adapted for *C. elegans* (Zhang et al., 2015). The degron-tagged CFI-1::wrmScarlet was generated by CRISPR/Cas9, and it was conditionally degraded in PVC neurons when exposed to auxin in the presence of PVC neuron-specific P*nmr-1*::TIR1. The indole-3-acetic acid (IAA) was dissolved in DMSO to prepare a 500 mM store solution, and it is preserved in a dark place at 4°C. The IAA solution was added to NGM (nematode growth medium) agar plates to a final concentration of 1 mM, and the plates could be shielded from light at 4°C for four weeks. Concentrated OP50 was seeded on the auxin NGM plates and allowed to dry overnight at room temperature. To induce CFI-1 degradation, L4 stage P*nmr-1*::TIR1; CFI-1::wrmScarlet::degron worms were transferred onto the IAA-coated plates and kept at 22°C. The worms were transferred to new IAA plates every 3 days. The CFI-1 expression pattern and UNC-9::GFP distribution were characterized in F1 young adults.

### Touch response analysis

Gentle touch was delivered with an eyelash affixed to a pipet tip. The posterior or the anterior part of the worm was touched with the eyelash edge in a top-down fashion. Young adult worms were placed on NGM plates with a thin layer of OP50 lawn. Responses to stimulation were recorded within 3**-**second window. Each worm was tested once. Each trial encompassed the evaluation of over 17 animals per genotype, with three independent replicates. All data are shown as mean ± s.d.

## Acknowledgments

We thank the Caenorhabditis Genetics Center (CGC, USA) and the National BioResource Project (NBRP, Japan) for providing strains. We also thank Dr. Cornelia I. Bargmann for providing the *cfi-1(ky650)* mutant strain.

## Author contributions

Conceptualization: Z.W., M.D.; Methodology: Z.W.; Formal analysis: Z.W., Investigation: Z.W.; L.P., Resources: Z.W., M.D; Data curation: Z.W.; Writing original draft: Z.W.; Writing - review & editing: Z.W., M.D., Supervision: M.D; Funding acquisition: M.D.

## Data availability

All relevant data can be found within the article and its supplementary information.

## Figure legends for supplemental figures

**Fig. S1. Gap junction distribution of P*unc-53* expressing neurons.** (A) Schematic drawing of the distribution of UNC-9::GFP puncta (green) formed by P*unc-53* expressing neurons. (B) Colocalization between P*unc-53*::UNC-9::GFP puncta and PVQ neuron (red) in the tail region. Scale bar: 25 µm. (C) P*unc-53*::UNC-9::GFP puncta in the head region in wild type (*wt*) and *cfi-1(xd424)* animals. (D) P*unc-53*::UNC-9::GFP puncta in the tail region in wild type (*wt*) and *cfi-1(xd424)* animals.

**Fig. S2. The endogenous expression of *cfi-1* gene.** (A and B) CFI-1::GFP knock-in (green) distribution in the head, ventral cord, and the tail region in L1 (A) and L2 (B) stage. Scale bar, 10 µm. (C) *cfi-1(xd424)* mutant phenotype could not be rescued by expressing wild type copy of *cfi-1* in the ventral cord region. Scale bar: 25 µm.

**Fig. S3. Identification of the P*unc-53* expressing neurons.** (A) *cfi-1* (red) (labeled by P*cfi-1*::myr::mCherry) is not expressed in Punc-53 expressing neurons (green). (B) P*lad-2* driven mCherry labels ALN and PLN neurons (red). (C) P*odr-2 2b* driven mCherry labels PVP neurons (red). (D) P*sra-6* driven mCherry labels PVQ neurons (red). (E) P*flp-7* driven mCherry labels PVW neurons (red). (F) P*flp-10* driven mCherry labels PQR and PVR neurons (red). (G) P*mec-7* driven mCherry labels ALN and PLM neurons (red). (H) P*pdf-1* driven mCherry labels PVN neurons (red). Scale bar: 10 µm.

**Fig. S4. The endogenous gap junction formation of DD/VD neurons is not affected by *cfi-1*.** (A and B) The UNC-9::split-GFP puncta distribution resembles P*unc-53*::UNC-9::GFP in the posterior ventral cord and tail region. (C) The distribution of endogenous gap junctions (labeled by P*unc-25*::UNC-9::split-GFP, green) in wild type (*wt*) and *cfi-1(xd424)* mutants. Scale bar: 25 µm. (D) Quantification of the P*unc-25*::UNC-9::split-GFP puncta number in wild type (*wt*) and *cfi-1(xd424)* mutants. Student’s *t*-test was performed. NS: not significant, N ≥ 50.

**Fig. S5. Neural circuitry underlying posterior touch response.** (A-C) The neural circuitry underlying posterior touch response in wild type (A), *cfi-1* mutant (B) with or without wild type *cfi-1* being expressed in PVR neuron (C). The touch cell connectors, LUA, and possible PVR, are marked by rectangles. The interneurons are marked by hexagons. The motor neurons are marked by circles. Both chemical synapses (→) and gap junctions (┤) are indicated. The PVC-PVR gap junction is highlighted (purple). The thickness of individual lines represents the different strength of signaling.

## References

Alcami, P. and Pereda A. E. (2019) Beyond plasticity: the dynamic impact of electrical synapses on neural circuits. Nat. Rev. Neurosci. 20, 253–271. doi: 10.1038/s41583-019-0133-5

Abascal, F. and Zardoya, R. (2013). Evolutionary analyses of gap junction protein families. Biochim. Biophys. Acta. 1828, 4–14. doi: 10.1016/j.bbamem.2012.02.007

An, G., Miner, C. A., Nixon, J. C., Kincade, P. W., Bryant, J., Tucker, P. W. and Webb, C. F. (2010). Loss of Bright/ARID3a function promotes developmental plasticity. Stem Cells 28, 1560–1567. doi: 10.1002/stem.491

Barnes, T. M., and Hekimi, S. (1997). The *Caenorhabditis elegans* avermectin resistance and anesthetic response gene *unc-9* encodes a member of a protein family implicated in electrical coupling of excitable cells. J Neurochem 69, 2251–2260. doi: 10.1046/j.1471-4159.1997.69062251.x

Barrios, A., Ghosh, R., Fang, C., Emmons, S.W. and Barr, M. M. (2012). PDF-1 neuropeptide signaling modulates a neural circuit for mate-searching behavior in *C. elegans*. Nat. Neurosci. 15, 1675–1682. doi: 10.1038/nn.3253

Berghoff, E. G., Glenwinkel, L., Bhattacharya, A., Sun, H., Varol, E., Mohammadi, N., Antone, A., Feng, Y., Nguyen, K., Cook, S.J. et al. (2021). The Prop1-like homeobox gene *unc-42* specifies the identity of synaptically connected neurons. eLife 10, e64903. doi: 10.7554/eLife.64903

Bhattacharya, A., Aghayeva, U., Berghoff, E.G. and Hobert, O. (2019). Plasticity of the electrical connectome of *C. elegans*. Cell 176, 1174–1189. doi: 10.1016/j.cell.2018.12.024

Brenner, S. (1974) The genetics of *Caenorhabditis elegans*. Genetics 77, 71–94. doi: 10.1016/j.ydbio.2022.02.013

Bruzzone, R. and Dermietzel, R. (2006). Structure and function of gap junctions in the developing brain. Cell Tissue Res. 326, 239–248. doi: 10.1007/s00441-006-0287-0

Chalfie, M., Sulston, J. E., White, J. G., Southgate, E., Thomson, J. N. and Brenner, S. (1985) The neural circuit for touch sensitivity in *Caenorhabditis elegans*. J. Neurosci. 5, 956–964. doi: 10.1523/JNEUROSCI.05-04-00956.1985

Choi. M. K., Liu, H., Wu, T., Yang, W. and Zhang, Y. (2020) NMDAR-mediated modulation of gap junction circuit regulates olfactory learning in *C. elegans*. Nat. Commun. 11, 3467. doi: 10.1038/s41467-020-17218-0

Chou, J.H., Bargmann, C.I. and Sengupta, P. (2001). The *Caenorhabditis elegans odr-2* gene encodes a novel Ly-6-related protein required for olfaction. Genetics 157, 211–224. doi: 10.1093/genetics/157.1.211

Connors, B. W. and Long, M. A. (2004). Electrical synapses in the mammalian brain. Annu. Rev. Neurosci. 27, 393–418. doi: 10.1146/annurev.neuro.26.041002.131128

Cook, S. J., Jarrell, T. A., Brittin, C. A., Wang, Y., Bloniarz, A. E., Yakovlev, M. A., Nguyen, K. C. Q., Tang, L. T. H., Bayer, E. A., Duerr, J. S. et al. (2019). Whole-animal connectomes of both *Caenorhabditis elegans* sexes. Nature 571, 63–71. doi: 10.1038/s41586-019-1352-7

Curtin, K.D., Zhang, Z. and Wyman, R.J. (2002) Gap junction proteins expressed during development are required for adult neural function in the *Drosophila* optic lamina. J. Neurosci. 22, 7088–7096. doi: 10.1523/JNEUROSCI.22-16-07088.2002

Dere, E. and Zlomuzica, A. (2012). The role of gap junctions in the brain in health and disease. Neurosci. Biobehav. Rev. 36, 206–217. doi: 10.1016/j.neubiorev.2011.05.015

Dykes, I. M., Freeman, F. M., Bacon, J. P. and Davies, J.A. (2004). Molecular basis of gap junctional communication in the CNS of the *Leech Hirudo* medicinalis. J. Neurosci. 24, 886–894. doi: 10.1523/JNEUROSCI.3676-03.2004

Elfgang, C., Eckert, R., Lichtenberg-Fraté, H., Butterweck, A., Traub, O., Klein, R.A., Hulser, D.F. and Willecke, K. (1995). Specific permeability and selective formation of gap junction channels in connexin-transfected HeLa cells. J. Cell Biol. 129, 805–817. doi: 10.1083/jcb.129.3.805

Evans, W. H. and Martin, P. E. (2002). Gap junctions: structure and function (Review). Mol Membr Biol 19, 121–136. doi: 10.1080/09687680210139839

Fukuda, T. (2017). Structural organization of the dendritic reticulum linked by gap junctions in layer 4 of the visual cortex. Neuroscience 340, 76–90. doi: 10.1016/j.neuroscience.2016.10.050

Gengyo-Ando, K. and Mitani, S. (2000) Characterization of mutations induced by ethylmethanesulfonate, UV, and trimethylpsoralen in the nematode *Caenorhabditis elegan*s. Biochem. Biophys. Res. Commun. 269, 64–69. doi: 10.1006/bbrc.2000.2260

Glenwinkel, L., Taylor, S. R., Langebeck-Jensen, K., Pereira, L., Reilly, M.B., Basavaraju, M., Rafi, I., Yemini, E., Pocock, R., Sestan, N. et al. (2021). In silico analysis of the transcriptional regulatory logic of neuronal identity specification throughout the *C. elegans* nervous system. eLife 10, e64906. doi: 10.7554/eLife.64906

Goodenough, D.. and Paul, D. L. (2009). Gap junctions. Cold Spring Harb. Perspect Biol. 1, a002576. doi: 10.1101/cshperspect.a002576

Greb, H., Hermann, S., Dirks, P., Ommen, G., Kretschmer, V., Schultz, K., Zoidl, G., Weiler, R. and Janssen-Bienhold, U. (2017). Complexity of gap junctions between horizontal cells of the carp retina. Neuroscience 340, 8–22. doi: 10.1016/j.neuroscience.2016.10.044

He, S., Cuentas-Condori, A. and Miller, D. M. 3rd. (2019) NATF (Native and Tissue-Specific Fluorescence): A strategy for bright, tissue-specific GFP labeling of native proteins in *Caenorhabditis elegans*. Genetics 212, 387–395. doi: 10.1534/genetics.119.302063

Herrscher, R. F., Kaplan, M. H., Lelsz, D. L., Das, C., Scheuermann, R. and Tucker, P. W. (1995) The immunoglobulin heavy-chain matrix-associating regions are bound by Bright: a B cell-specific trans-activator that describes a new DNA-binding protein family. Genes Dev. 9, 3067–3082. doi: 10.1101/gad.9.24.3067

Hurkey, S., Niemeyer, N., Schleimer, J.H., Ryglewski, S., Schreiber, S. and Duch, C. (2023). Gap junctions desynchronize a neural circuit to stabilize insect flight. Nature 618, 118–125. doi: 10.1038/s41586-023-06099-0

Jabeen, S. and Thirumalai, V. (2018). The interplay between electrical and chemical synaptogenesis. J. Neurophysiol. 120, 1914–1922. doi: 10.1152/jn.00398.2018

Joseph, S., Deneke, V. E. and Cowden Dahl, K. D. (2012) ARID3B induces tumor necrosis factor alpha mediated apoptosis while a novel ARID3B splice form does not induce cell death. PLoS One 7, e42159. doi: 10.1371/journal.pone.0042159

Kerk, S. Y., Kratsios, P., Hart, M., Mourao, R. and Hobert, O. (2017) Diversification of *C. elegans* motor neuron identity via selective effector gene repression. Neuron 93, 80–98. doi: 10.1016/j.neuron.2016.11.036

Kim, K. and Li, C. (2004). Expression and regulation of an FMRFamide-related neuropeptide gene family in *Caenorhabditis elegans*. J. Comp. Neurol. 475, 540–550. doi: 10.1002/cne.20189

Kortschak, R. D., Tucker, P. W. and Saint, R. (2000). ARID proteins come in from the desert. Trends Biochem. Sci. 25, 294–299. doi: 10.1016/s0968-0004(00)01597-8

Kratsios, P., Kerk, S. Y., Catela, C., Liang, J., Vidal, B., Bayer, E. A., Feng, W., De La Cruz, E.D., Croci, L., Consalez, G.G. et al. (2017). An intersectional gene regulatory strategy defines subclass diversity of *C. elegans* motor neurons. eLife 6. e25751. doi: 10.7554/eLife.25751

Li, Y., Smith, J. J., Marques, F., Osuma, A., Huang, H. C. and Kratsios, P. (2023). Cell context-dependent CFI-1/ARID3 functions control neuronal terminal differentiation. Cell Rep. 42, 112220. doi: 10.1016/j.celrep.2023.112220

Lopresti, V., Macagno, E. R. and Levinthal, C. (1974) Structure and development of neuronal connections in isogenic organisms: transient gap junctions between growing optic axons and lamina neuroblasts. Proc. Natl. Acad. Sci. USA 71, 1098–1102. doi: 10.1073/pnas.71.4.1098

Ma, X., Zhao, Z., Xiao, L., Xu, W., Kou, Y., Zhang, Y., Wu, G., Wang, Y. and Du, Z. (2021). A 4D single-cell protein atlas of transcription factors delineates spatiotemporal patterning during embryogenesis. Nat. Methods 18, 893–902. doi: 10.1038/s41592-021-01216-1

Martin, E. A., Michel, J. C., Kissinger, J. S., Echeverry, F. A., Lin, Y. P., O’Brien, J., Pereda, A.E. and Miller, A. C. (2023). Neurobeachin controls the asymmetric subcellular distribution of electrical synapse proteins. Curr. Biol. 33, 2063–2074. doi: 10.1016/j.cub.2023.04.049

Meng, L., and Yan, D. (2020). NLR-1/CASPR Anchors F-Actin to Promote Gap Junction Formation. Dev. Cell 55, 574–587 e573. doi: 10.1016/j.devcel.2020.10.020

Miller, A. C., Voelker, L. H., Shah, A. N. and Moens, C. B. (2015). Neurobeachin is required postsynaptically for electrical and chemical synapse formation. Curr. Biol. 25, 16–28. doi: 10.1016/j.cub.2014.10.071

Mitani, S., Du, H., Hall, D. H., Driscoll, M. and Chalfie, M. (1993) Combinatorial control of touch receptor neuron expression in *Caenorhabditis elegans*. Development 119, 773–783. doi: 10.1242/dev.119.3.773

Numata, S., Claudio, P. P., Dean, C., Giordano, A. and Croce, C. M. (1999) Bdp, a new member of a family of DNA-binding proteins, associates with the retinoblastoma gene product. Cancer Res. 59, 3741–3747.

O’Rourke, N. A., Weiler, N. C., Micheva, K. D. and Smith, S. J. (2012). Deep molecular diversity of mammalian synapses: why it matters and how to measure it. Nat. Rev. Neurosci. 13, 365–379. doi: 10.1038/nrn3170

Oshima, A., Matsuzawa, T., Murata, K., Tani, K. and Fujiyoshi, Y. (2016) Hexadecameric structure of an invertebrate gap junction channel. J. Mol. Biol. 428, 1227–1236. doi: 10.1016/j.jmb.2016.02.011

Palumbos, S. D., Skelton, R., McWhirter, R., Mitchell, A., Swann, I., Heifner, S., Von Stetina, S. and Miller, D. M. 3rd. (2021) cAMP controls a trafficking mechanism that maintains the neuron specificity and subcellular placement of electrical synapses. Dev. Cell 56, 3235-3249. doi: 10.1016/j.devcel.2021.10.011

Pereda, A.E. (2014). Electrical synapses and their functional interactions with chemical synapses. Nat. Rev. Neurosci. 15, 250–263. doi: 10.1038/nrn3708

Phelan, P., Stebbings, L. A., Baines, R. A., Bacon, J. P., Davies, J. A. and Ford, C. (1998). Drosophila shaking-B protein forms gap junctions in paired *Xenopus* oocytes. Nature 391, 181–184. doi: 10.1038/34426

Rabinowitch, I., Chatzigeorgiou, M., Zhao, B., Treinin, M. and Schafer, W.R. (2014). Rewiring neural circuits by the insertion of ectopic electrical synapses in transgenic *C. elegans*. Nat. Commun. 5, 4442. doi: 10.1038/ncomms5442

Rash, J.E., Curti, S., Vanderpool, K. G., Kamasawa, N., Nannapaneni, S., Palacios-Prado, N., Flores, C.E., Yasumura, T., O’Brien, J., Lynn, B.D. et al. (2013). Molecular and functional asymmetry at a vertebrate electrical synapse. Neuron 79, 957–969. doi: 10.1016/j.neuron.2013.06.037

Ratliff, M.L., Mishra, M., Frank, M.B., Guthridge, J.M., and Webb, C.F. (2016). The transcription factor ARID3a is important for *in vitro* differentiation of human hematopoietic progenitors. J. Immunol. 196, 614–623. doi: 10.4049/jimmunol.1500355

Ratliff, M.L., Templeton, T.D., Ward, J.M., and Webb, C.F. (2014). The bright side of hematopoiesis: regulatory roles of ARID3a/Bright in human and mouse hematopoiesis. Front Immunol. 5, 113. doi: 10.3389/fimmu.2014.00113

Roy L, Samyesudhas SJ, Carrasco M, Li J, Joseph S, Dahl R, Cowden Dahl KD (2014) ARID3B increases ovarian tumor burden and is associated with a cancer stem cell gene signature. Oncotarget 5, 8355–8366. doi: 10.18632/oncotarget.2247

Shaham S, Bargmann CI (2002) Control of neuronal subtype identity by the *C. elegans* ARID protein CFI-1. Genes Dev. 16, 972–983. doi: 10.1101/gad.976002

Shandala, T. et al. (1999) The *Drosophila* dead ringer gene is required for early embryonic patterning through regulation of argos and button head expression. Development 126, 4341–4349. doi: 10.1242/dev.126.19.4341

Söhl G, Maxeiner S, Willecke K (2005) Expression and functions of neuronal gap junctions. Nat. Rev. Neurosci. 6, 191–200. doi: 10.1038/nrn1627

Sosinsky, G.E. and Nicholson, B.J. (2005). Structural organization of gap junction channels. Biochim Biophys Acta 1711, 99–125. doi: 10.1016/j.bbamem.2005.04.001

Stauffer KA, Kumar NM, Gilula NB, Unwin N (1991) Isolation and purification of gap junction channels. J. Cell Biol. 115, 141-150. doi: 10.1083/jcb.115.1.141

Stringham E, Pujol N, Vandekerckhove J, Bogaert T (2002) *unc-53* controls longitudinal migration in *C. elegans*. Development 129, 3367–3379. doi: 10.1242/dev.129.14.3367

Südhof, T.C. (2012). The presynaptic active zone. Neuron 75, 11–25. doi: 10.1016/j.neuron.2012.06.012

Talukdar, S., Emdad, L., Das, S. K. and Fisher, P. B. (2022). GAP junctions: multifaceted regulators of neuronal differentiation. Tissue Barriers 10, 1982349. doi: 10.1080/21688370.2021.1982349

Teubner, B., Degen, J., Söhl, G., Guldenagel, M., Bukauskas, F. F., Trexler, E.B., Verselis, V. K., De Zeeuw, C.I., Lee, C.G., Kozak, C.A. et al. (2000). Functional expression of the murine connexin 36 gene coding for a neuronspecific gap junctional protein. J. Membr. Biol. 176, 249–262. doi: 10.1007/s00232001094

Troemel, E. R., Chou, J. H., Dwyer, N. D., Colbert, H. A. and Bargmann, C. I. (1995). Divergent seven transmembrane receptors are candidate chemosensory receptors in *C. elegans*. Cell 83, 207–218. doi: 10.1016/0092-8674(95)90162-0

Valentine, S. A., Chen, G., Shandala, T., Fernandez, J., Mische, S., Saint, R. and Courey, A. J. (1998). Dorsal-mediated repression requires the formation of a multiprotein repression complex at the ventral silencer. Mol. Cell Biol. 18, 6584–6594. doi: 10.1128/MCB.18.11.6584

Vervaeke, K., Lorincz, A., Gleeson, P., Farinella, M., Nusser, Z. and Silver, R. A. (2010) Rapid desynchronization of an electrically coupled interneuron network with sparse excitatory synaptic input. Neuron 67, 435–451. doi: 10.1016/j.neuron.2010.06.028

Walker, D. S. and Schafer, W. R. (2020). Distinct roles for innexin gap junctions and hemichannels in mechanosensation. eLife 9, e50597. doi: 10.7554/eLife.50597

Wang, X., Zhang, W., Cheever, T., Schwarz, V., Opperman, K., Hutter, H., Koepp, D. and Chen, L. (2008). The *C. elegans* L1CAM homologue LAD-2 functions as a coreceptor in MAB-20/Sema2 mediated axon guidance. J. Cell Biol. 180, 233–246. doi: 10.1083/jcb.200704178

White, J. G., Southgate, E., Thomson, J. N. and Brenner, S. (1986) The structure of the nervous system of *Caenorhabditis elegans*. Philos. Trans. R. Soc. Lond. B. Biol. Sci. 314, 1–340. doi: 10.1098/rstb.1986.0056

White, J. G., Southgate, E. and Thomson, J. N. (1992). Mutations in the *Caenorhabditis elegans unc-4* gene alter the synaptic input to ventral cord motor neurons. Nature 355, 838–841. doi: 10.1038/355838a0

Wilsker, D., Probst, L., Wain, H.M., Maltais, L., Tucker, P.W. and Moran, E. (2005). Nomenclature of the ARID family of DNA-binding proteins. Genomics 86, 242–251. doi: 10.1016/j.ygeno.2005.03.013

Webb, C.F., Bryant, J., Popowski, M., Allred, L., Kim, D., Harriss, J., Schmidt, C., Miner, C.A., Rose, K., Cheng, H.L. et al. (2011). The ARID family transcription factor bright is required for both hematopoietic stem cell and B lineage development. Mol. Cell Biol. 31, 1041–1053. doi: 10.1128/MCB.01448-10

Yao, X.H., Wang, M., He, X. N., He, F., Zhang, S. Q., Lu, W., Qiu, Z. L. and Yu, Y. C. (2016). Electrical coupling regulates layer 1 interneuron microcircuit formation in the neocortex. Nat. Commun. 7, 12229. doi: 10.1038/ncomms12229

Yeh, E., Kawano, T., Ng, S., Fetter, R., Hung, W., Wang, Y. and Zhen, M. (2009) *Caenorhabditis elegans* innexins regulate active zone differentiation. J. Neurosci. 29, 5207–5217. doi: 10.1523/JNEUROSCI.0637-09.2009

Zhang, L., Ward, J. D., Cheng, Z. and Dernburg, A. F. (2015) The auxin-inducible degradation (AID) system enables versatile conditional protein depletion in *C. elegans*. Development 142, 4374–4384. doi: 10.1242/dev.129635

Zolnik, T. A. and Connors, B. W. (2016). Electrical synapses and the development of inhibitory circuits in the thalamus. J. Physiol. 594, 2579–2592. doi: 10.1113/JP271880

Zhang, J., Li, X., Jevince, A. R., Guan, L., Wang, J., Hall, D. H., Huang, X. and Ding, M. (2013). Neuronal target identification requires AHA-1-mediated fine-tuning of Wnt signaling in *C. elegans*. PLoS Genet 9, e1003618. doi: 10.1371/journal.pgen.1003618

